# High-resolution temporal transcript profiling during *Arabidopsis thaliana* gynoecium morphogenesis uncovers the chronology of gene regulatory network activity and reveals novel developmental regulators

**DOI:** 10.1101/2020.07.29.227314

**Authors:** Kimmo I. Kivivirta, Denise Herbert, Clemens Roessner, Stefan de Folter, Nayelli Marsch-Martinez, Annette Becker

## Abstract

The gynoecium is the most complex organ formed by the flowering plants. It encloses the ovules, provides a surface for pollen contact and self-incompatibility reactions, allows pollen tube growth and, post fertilization, and develops into the fruit. Consequently, the regulation of gynoecium morphogenesis is complex and appropriate timing of this process in part determines reproductive success. However, little is known about the global control of gynoecium development, even though many regulatory genes have been characterized. Here, we characterized dynamic gene expression changes using laser-microdissected gynoecium tissue from four developmental stages in Arabidopsis. We provide a high-resolution map of global expression dynamics during gynoecium morphogenesis and link these to the gynoecium interactome. We reveal groups of genes acting together early and others acting late in morphogenesis. Clustering of co-expressed genes enables comparisons between the leaf, shoot apex, and gynoecium transcriptomes allowing the dissection of common and distinct regulators. Furthermore, our results lead to the discovery of the *LESSER FERTILITY1-4* (*LEF1-4*) genes, which, when mutated, lead to impaired gynoecium expansion, illustrating that global transcriptome analyses reveal yet unknown developmental regulators. Our data show that highly interacting proteins, such as *SEPALLATA3, AGAMOUS*, and *TOPLESS* are expressed more evenly during development, but switch interactors in time, whereas stage-specific proteins have only few interactors. Our analysis connects specific transcriptional regulator activities, protein interactions, and underlying metabolic processes towards the development of a dynamic network model for gynoecium development.

## Introduction

Broadening our understanding of flower development is important as most of the terrestrial life is either directly or indirectly dependent on flowering plants (Sauquet et al., 2017) and agricultural advancements are required to feed the growing global population of the 21st century. Carpels, the female reproductive organs of the flowering plants begin to develop after the plant has reached its generative maturity and flowering has initiated. Carpels are located in the innermost whorl of the flower and their sum is defined as gynoecium. The gynoecium bears the developing ovules, receives pollen grains, and allows their passage through specialized tissue to enable fertilization of the ovules. Subsequently, these develop into seeds while the gynoecium is converted into a fruit.

In *Arabidopsis thaliana*, flowers arise on the flanks of the inflorescence meristem. The flower consists of four concentric whorls of different organs: the outermost sepals, then follow petals, stamina, and the gynoecium is formed in the centre. Gynoecium development commences approximately four days after floral development initiation when the previously undifferentiated central dome in the middle of the flower starts to elongate and forms a hollow, oval shape. This tube-like gynoecium consists of two congenitally fused carpels (Smyth et al., 1990). Inside the gynoecium, the carpel margin meristem (CMM) initiates as the inner adaxial margins first bulge inward forming a boundary surface inside the hollow structure. The CMM then gives rise to the carpel marginal tissues from where placenta, ovules, false septum, and transmitting tract form (Bowman et al., 1999, Reyes-Olalde et al., 2013, Reyes-Olalde and de Folter, 2019). The septa primordia fuse and form the false septum through postgenital fusion. After approximately eleven days of flower development, stigmatic papillae start to appear at the tip of the developing organ. One day later the papillae fully cover the tip of the gynoecium and the open-ended structure closes by postgenital fusion, while style and transmitting tract differentiate, leading to the mature gynoecium (Smyth et al., 1990).

The initiation of the gynoecium requires activation of the class C and E homeotic genes *AGAMOUS (AG)* and *SEPALLATA3 (SEP3)* (Bowman et al., 1989, Honma and Goto, 2001, Pelaz et al., 2000). These proteins form a tetramer protein complex with the active sites binding to a plethora of promoter regions in the Arabidopsis genome regulating the expression of the downstream genes to provide carpel organ identity and initiate carpel development (Smaczniak et al., 2017).

Post initiation, the dome-shaped floral meristem differentiates into several tissue types. These require specification and orientation towards the adaxial/abaxial and apical/basal axes, processes controlled by transcriptional regulators (TRs) such as *PHABULOSA (PHB), REVOLUTA (REV), PHAVOLUTA (PHV), NUBBIN (NUB), JAGGED (JAG)* and others (McConnell et al., 2001, Bowman et al., 2002, Dinneny et al., 2006). Induction and differentiation of the CMM tissues is regulated by *SPATULA (SPT), CUP-SHAPED COTYLEDON1-2 (CUC1-2), HECATE1-3 (HEC1-3), INDEHISCENT (IND)* for example (Heisler et al., 2001, Aida & Tasaka 2006, Gremski et al., 2007, Kay et al., 2012) and differentiation of stigma and style by *NGATHA3 (NGA3), STY1, STY2* etc. (Trigueros et al., 2009, Sessions & Zambryski 1995, Kuusk et al., 2002). A complex interplay of many additional genes, phytohormones, peptides, microRNAs and epigenetic factors ultimately lead to the complete organogenesis of the gynoecium (reviewed in further detail in Alvarez-Buylla et al., 2010, Krishnamurthy & Bahadur 2015 and Moubayidin & østergaard 2017).

While genetic and protein interactions of many of the TRs coordinating carpel development are known (reviewed in Reyes-Olalde et al., 2013, Chávez Montes et al., 2015, Zúñiga-Mayo et al., 2019, and Becker 2019), we lack a comprehensive picture of expression dynamics of these TRs during carpel development. So far, the major transcriptomic studies of flower development in *A. thaliana* have focused on either the later stages of the developed flower organs (Klepikova et al., 2016) or complete buds at early to late stages (Ryan et al., 2015 and Mantegazza et al., 2014). Here, we provide a high-resolution temporal transcription time scale map of gynoecium development in *A. thaliana*, based on laser-microdissection (LMD) with subsequent RNAseq analysis of four different stages of carpel development starting from the initiation of carpel development to maturation, excluding the ovules. We show that specific genetic modules exist in a temporally precisely regulated manner and identify consecutively acting protein interaction networks key to gynoecium development. Further, we identify four putative transcription factors (*LESSER FERTILITY1-4, LEF1-4*) based on their specific temporal expression during gynoecium development and show that they contribute to gynoecium longitudinal growth and seed formation.

## Results

### Arabidopsis transcriptome data of four stages of carpel development

We sequenced laser-microdissected Arabidopsis carpel RNA samples at four different developmental stages: S1, initiation of carpel development after the differentiation of the central dome corresponding to stage 5 of *A. thaliana* flower development (Smyth et al., 1990); S2, elongation of carpel walls (stage 9); S3, during the female meiosis (stage 11, Smyth et al., 1990; Armstrong & Jones, 2001); S4, between female meiosis and anthesis (stage 12). Sample preparation, RNA-seq, transcriptome assembly and quality control are described elsewhere (Kivivirta et al., 2019). Four biological replicates were sequenced for all the four developmental stages and three were used for this analysis. 33 Mio paired-end reads were sequenced with read length of approximately 76 bp and annotated, resulting in expression information of all *A. thaliana* genes during gynoecium development.

### Expression dynamics of carpel developmental regulators

We were interested in the temporal expression profiles of known carpel developmental genes to learn if the timing of their expression matches with their known role in development. We analysed carpel regulatory genes by generating an expression heatmap (Fig. 1, Supplemental table 1).

**Figure 1:**
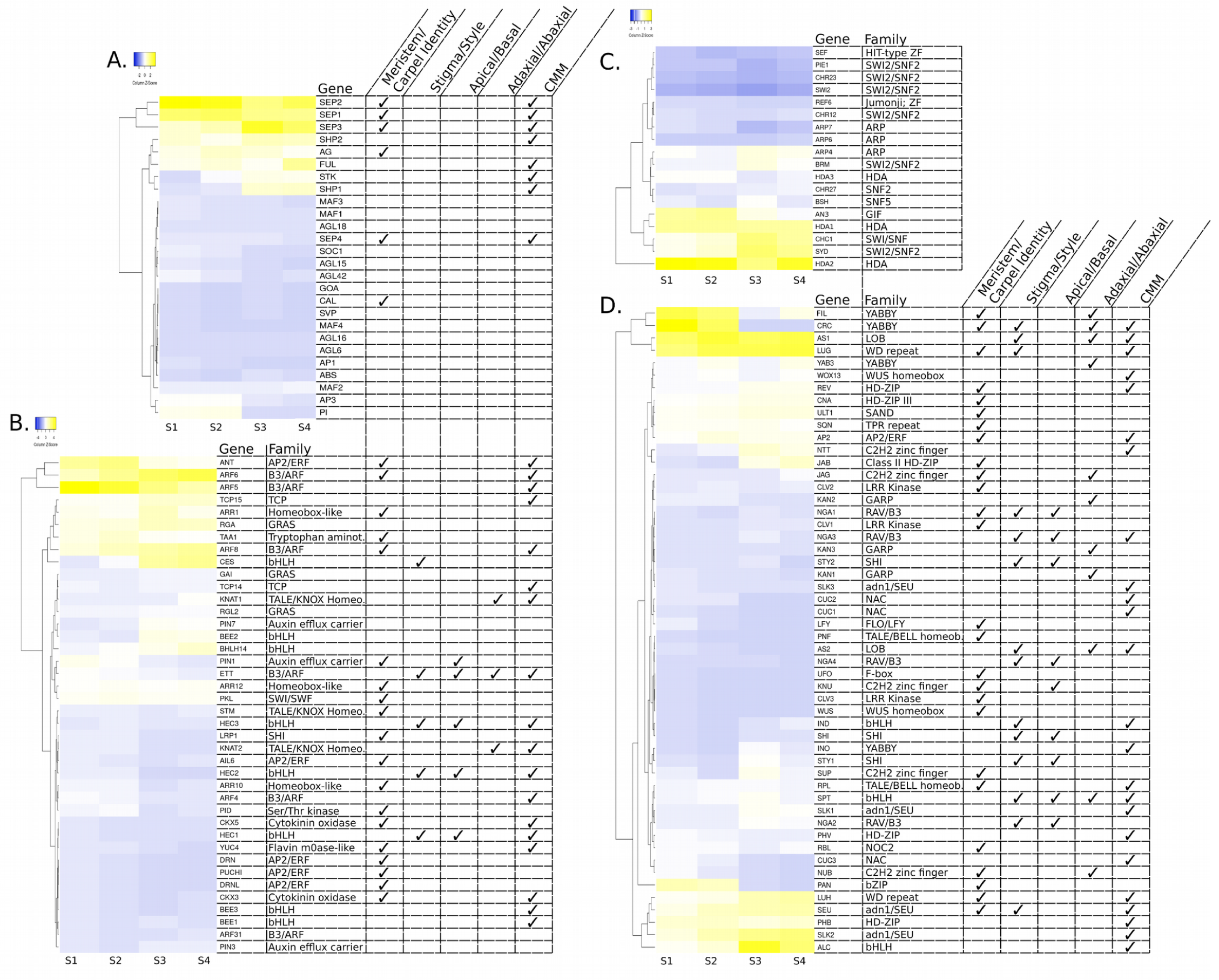
The carpel regulome heatmap. Heatmap of carpel developmental genes illustrating the strength of gene expression during four developmental stages, strongly expressed genes are bright yellow, weakly expressed genes are in dark blue. Similarly expressed genes were clustered with Euclidean distance for the absolute values of gene expression. A) Heatmap of MIKC type MADS-box genes transcriptionally active in the carpel (the full set is available in Supplemental Figure 1). B) Genes involved in phytohormone signalling, homeostasis, perception, or biosynthesis, or related to phytohormone pathways, with known regulatory functions during carpel development. C) Genes involved in chromatin remodeling. Functions are not shown for the chromatin remodelers as their role in gynoecium development is unclear. D) A collection of other regulatory genes required for carpel development. The table to the right (A, B, D) of the heat map indicates the gene’s contribution to carpel developmental regulation: organ identity, development of stigma/style, apical/basal and adaxial/abaxial patterning and CMM development (references for gene functions in Supplemental Table 1) and, in B), C) and D) their gene family membership. Genes with average expression of TPM <1 were omitted from the analysis.

Among the genes most important for floral organ identity, initiation and maintenance are the MIKC MADS-box transcription factors *SEP1-4* and *AG* (Fig. 1A, Supplemental figure 1). While SEP1-4 show strong differences in expression dynamics, *AG* is expressed evenly at a low level throughout the stages. The *APETALA3* (*AP3*) and *PISTILLATA* (*PI*) genes required for stamen and petal but not gynoecium organ identity, show expression in the first two stages of gynoecium development, confirming earlier observations (Goto and Meyerowitz, 1994). Interestingly, some late acting MADS-box genes required for fruit dehiscence, such *FRUITFULL* (*FUL*) and *SHATTERPROOF2* (*SHP2*) are expressed strongly throughout gynoecium development.

Hormonal signalling is an integral part of carpel development, with crucial functions for signalling pathways such as auxin and cytokinin, but also others like brassinosteroids and gibberellins (Marsch-Martinez and de Folter, 2016; Zúñiga-Mayo et al., 2019). We observed expression of many of the genes and transcription factors related to these hormonal pathways (Fig. 1B). The genes that present the highest expression are involved in auxin and cytokinin regulation and response, auxin biosynthesis, and brassinosteroid regulation.

Genes related to different steps in the auxin pathway were identified, such as those coding for TAA and YUC (biosynthesis); PIN1, PIN3 and PIN7 (transport); PID (transporter regulation); and the response factors ARF5/MONOPTEROS, ARF3/ETTIN, ARF6, ARF8 and others. Also, transcription factors such as ANT and AIL6, among others, which are closely related to the auxin pathway, were found in the transcriptome data (Krizek, 2009). Moreover, we observed various transcription factors well known for their regulatory role in carpel development (Fig. 1D) that also affect auxin signalling such as *STY1, STY2, NGA, SPT*, and *CRABS CLAW* (*CRC*), or that respond to auxin such as the *CUC1-3* genes.

Genes related to the cytokinin pathway include those encoding the response regulators ARR1, ARR10 and ARR12, and the cytokinin degradation enzymes CKX3 and CKX5. All these genes have been reported to be expressed during gynoecium development, particularly in meristematic tissues. Mutations in these genes cause reduced or increased meristematic activity, respectively (Reyes-Olalde et al. 2017; Bartrina et al., 2011). Also, transcription factors such as the KNOX family members STM, BP and KNAT2, and TCP14 and TCP15 were expressed at different stages, play important roles in gynoecium development, and have been associated to the cytokinin pathway (Lucero et al., 2015).

Brassinosteroids also play important roles in gynoecium development. In the transcriptome data, the brassinosteroid-related genes *HALF FILLED (HAF/CES), BEE1, BEE2* and *BEE3*, were also expressed, specially at the intermediate and late stages of development. This is in line with their function in transmitting tract development later during gynoecium development (Crawford et al., 2011).

Gibberellins have recently been implicated in the negative modulation of ovule number (Gomez et al., 2018; Barro-Trastoy et al., 2020). DELLA proteins are negative regulators of gibberellin signalling, and their activity correlates positively with ovule number. Genes encoding for DELLA proteins such as *GAI, RGA, RGL2* were also found in the transcriptomes. Of these, only *RGA* is strongly expressed in the later stages, while the other show mild expression, decreasing in time.

Some of the genes in the transcriptomes take part in networks that connect different pathways. For example, the *HEC1-3* induce auxin signalling and repress cytokinin signalling in the style (Schuster et al., 2015). Another example is *SPT*, that besides inducing auxin biosynthesis and transport, activates the cytokinin response regulator *ARR1*, which in turn, also activates auxin biosynthesis and transport (Reyes-Olalde et al., 2017).

Chromatin remodelling is an essential component of plant development (Ojolo et al., 2018) but its involvement in gynoecium development has received little attention and we were interested in exploring whether known chromatin remodelers are differentially expressed during gynoecium development. *HISTONE DEACETYLASES1* and *2* (*HDA1/2*) are strongly expressed during gynoecium development whereas *HDA3* shows only little expression (Fig. 1C*). ACTIN-RELATED PROTEIN4* (*ARP4*), *BRAHMA* (*BRM*), *SPLAYED* (*SYD*), and *CHC1/SWP73B* show expression largely restricted to the latter two stages. In contrast, *GIF1/AN3’s* expression is mainly confined to the two early.

Several other TRs, not members of MADS-box genes, chromatin remodelers, or phytohormone-associated genes contribute essential functions to carpel morphogenesis (Fig. 1D). Among those, *CRC, FILAMENTOUS FLOWER (FIL), AS1, LUG, SEUSS (SEU), SEUSS-like2 (SLK2)*, and *LEUNIG*-*HOMOLOG* (*LUH*), *PHB* and *ALC* are most strongly expressed. In contrast, many other important regulators, such as *CUC1, CUC2*, or *WUS* are expressed at very low levels suggesting that even genes expressed at low level may profound impact on gynoecium development.

In summary, our high-resolution data confirm previously reported expression data for individual genes and shows differentiation of expression of regulatory genes, even between closely related homologs, such as *SHP1* and *SHP2* or the *SEP1-4* genes. Moreover, we can now identify temporal changes in regulatory gene activation during gynoecium development.

### Temporal dynamics of protein interactions

Transcriptional regulators often interact in dimers or higher order multimers, and for *A. thaliana* gynoecium development, many protein interactions of TRs have been identified. However, we were interested in the temporal dynamics of these protein interactions. Thus, a comprehensive carpel protein interactome was generated based on protein interactions previously verified by Yeast Two-Hybrid (Y2H), Bimolecular Fluorescence Complementation (BiFC) and/or Co-Immunoprecipitation (Co-IP) analyses (Fig. 2, Supplemental table 2). We overlaid this interaction with expression data to illustrate the transient nature of some gynoecium TR interactions.

**Figure 2:**
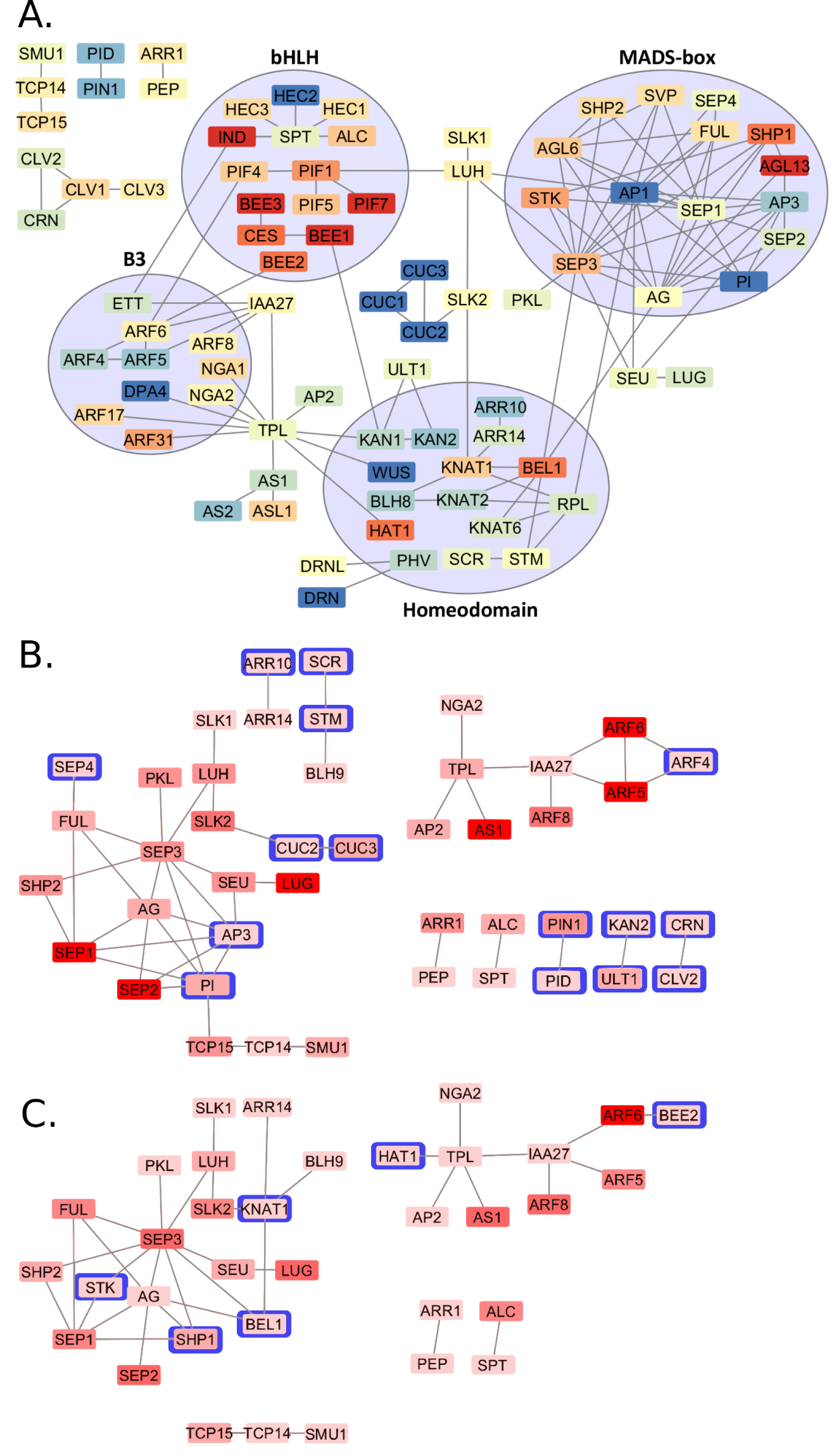
Temporal dynamics of the carpel regulatory interactome. A) Interaction map illustrating protein-protein interactions of important carpel developmental regulators based on experimentally verified interactions (Franz et al., 2018). An expression map (A) with the node colour indicating the trend of expression (logarithmic change of expression values) through the four carpel developmental stages (blue – yellow - red): blue node colour indicates a decreasing expression, yellow a stable expression, and red an increase in expressional strength through the four stages. Circles indicate membership of interacting proteins to larger transcription factor families. B) and C) Expression data showing only highly expressed interactions (> TMP 20) in developmental stage S1 (B) and S4 (C). The colour coding indicates the strength of expression, darker red nodes indicate high expression, light red nodes weak expression. The nodes with a blue frame show protein interaction partners unique to the respective stage. Proteins without verified interactions with TRs related to gynoecium development were omitted from the interaction maps.

Fig. 2A shows the contribution of single proteins and TR families to the carpel interactome. A group of several MADS-box proteins forms a highly interactive cluster, as do the bHLH, B3, and homeodomain transcription factor families. These families show different levels of connectivity among each other and with regulators outside of their family: The MADS-box proteins are highly connected to each other but interact with only five unrelated proteins. In contrast, the homeodomain proteins are less connected within their family, but interact with nine proteins outside their family.

Several hub proteins with five or more interactions were identified from the network analysis (Fig. 2A): the bHLH protein SPT, the B3 AUXIN RESPONSE FACTOR6 (ARF6), the transcriptional repressor INDOLE-3-ACETIC ACID INDUCIBLE 27 (IAA27) of the AUXIN/INDOLE-3-ACETIC ACID protein family, the WD40 transcriptional corepressor TOPLESS (TPL), and the homeodomain proteins BELL1 (BEL1), KNAT1/BREVIPEDICELLUS, REPLUMLESS (RPL), and BEL1-LIKE HOMEODOMAIN9 (BLH9). Moreover, the MADS-box proteins AG, PI, AP3, SEP1, SEP2, SEP3, AGAMOUS-LIKE6 (AGL6), FUL, SHP1, APETALA1 (AP1) act as hubs. Interestingly, the majority of hub protein encoding genes (*BLH9, TPL, SPT, AGL6, AP1, SEP1, AG*, and *AP3*) are expressed strongly (>TPM20) throughout all carpel developmental stages (Supplemental table 1). Only PI is expressed strongly only in early developmental stages (Fig. 2B), and also two are strongly expressed in late developmental stages (*SHP1* and *SEP3*) (Fig. 2C).

Many of the interacting hub proteins are generally rather strongly expressed (e.g. *SEP3* with TPM peak 297, *LUG* with 252, and *SEU* with 121), but their dynamics and interactivity change during development (Fig. 2B and C). The MADS box proteins *SEP4* and *PI* especially are highly expressed in S1 but when the gynoecium matures, their expression is reduced and other genes encoding highly interacting proteins like SEP3, SHP1, SHP2, STK, and FUL show increased expression.

Interestingly, not only hub genes, but also proteins with few interactors show stable expression throughout carpel development (Fig. 2B, C). The expression of each member in the cluster of the interacting proteins NGA2-TPL-AP2-AS1-IAA27-ARF5-ARF6-ARF8 remains remarkably stable (Fig. 2A, B). However, this cluster is complemented by the interaction of ARF4 with ARF5 and ARF6 at S1 which is not found in S4. Conversely, S4, interactions of HAT1/JAIBA with TPL and BEE2 with ARF6 are established.

In addition, the networks of TCP15-TCP14-SMU1, SLK1-LUH-SLK2, and LUG-SEU-SEP3-PKL-FUL-SHP2-SEP1-AG are stable throughout carpel development. Interestingly, other proteins also supplement these stable networks in different stages: The TCP15-TCP14-SMU1 network connects to PI in S1 and disconnects from the MADS-box protein cluster in S4. The SLK1-LUH-SLK2 network is connected to CUC2-CUC3 in S1 and exchanges this connection with KNAT1 in S4. The LUG-SEU-SEP3-PKL-FUL-SHP2-SEP1-SEP2-AG network is modified by the addition of SEP4, AP3, and PI in S1 and by STK, SHP1 and BEL1 in S4.

The interactome of S1 of carpel development includes 15 stage specific proteins while the S4 interactome includes only six stage specific proteins, and 29 proteins are included in the interactomes of both stages. This suggests that initiation and early morphogenesis of the carpel require more TR interactions than the later stages, when tissue differentiation is completed.

Another interesting group of proteins includes hub proteins (Fig. 2A) that have interaction partners with a generally or temporally very low level of expression. For example, BEL1 interacts with KNAT2 and KNAT6, but their expression is at a low level and at different stages, such that chances are high that the proteins never meet *in planta*. The same may apply to interactions with AGL6 and AP1, as the former has six and the latter thirteen protein interactors but they are hardly expressed in the gynoecium (Fig. 1A and 2A). Similar scenarios apply to KNAT1, SPT, and IAA27, which are all expressed at a low level. An extreme example is RPL, which has six interaction partners but is expressed at a very low level throughout gynoecium development and may be active mainly after fertilization during fruit development.

In summary, protein interactions directing carpel morphogenesis are temporally very dynamic. Only a few hub proteins maintain a high number of interactions throughout carpel development, such as SEP3, AG, and SEP1. Some components of the network, such as the one centred on TPL is active throughout carpel development but changes few interacting partners during morphogenesis. Further, differences in connectivity between transcription factor families were observed: while MADS-box proteins mainly interact among themselves, the bHLH family proteins are also highly connected with members of other TF families.

### Co-expression analyses provide comprehensive information on expression patterns and resulting shifts in biological processes

We were then interested in identifying genes that were co-regulated with the previously described carpel regulators to identify clusters of co-expressed and possibly co-regulated genes. Further, we aimed to learn if the carpel transcriptomes share more similarity with the leaf or SAM transcriptomes. Automatically partitioned clusters were generated to visualize co-expressed genes (Fig. 3, for the full list of clusters and genes see Supplemental table 3) within the four carpel development stages in comparison to leaf and SAM.

**Figure 3:**
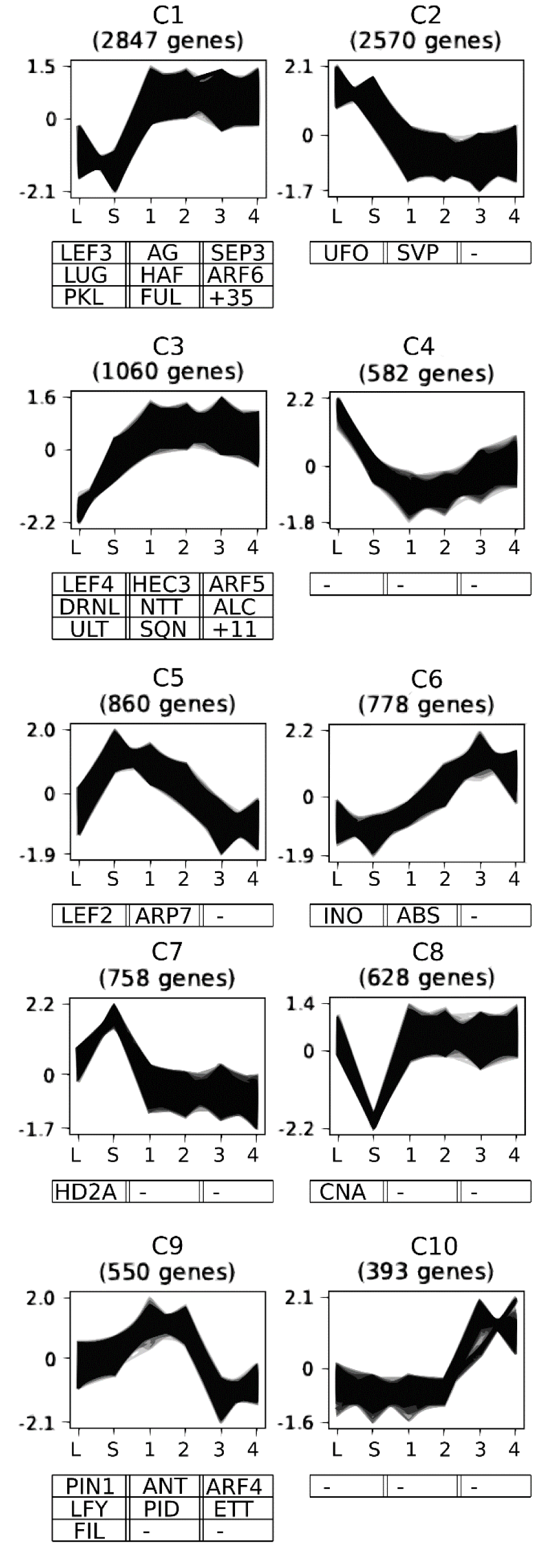
Clusters of co-expressed genes during gynoecium development. The strength of expression (Y axis) is illustrated for the different tissues and developmental stages of gynoecium development (X axis). The transcriptome files include the leaf blade (L), SAM (S) and four stages of gynoecium development (S1, S2, S3, S4). Example of known regulators are shown below each cluster. For the complete list of genes and GO analyses see Supplemental figure 3.

The largest cluster consists of genes exclusively upregulated throughout gynoecium development (C1: 2847 genes) and includes several well-known gynoecium developmental regulators such as *AG, SEP1-4, SHP1* and *2, SEU, SLK1-3, CES, LUG, LUH, HAF*, or *FUL*. The second-largest cluster (C2: 2570 genes) includes genes that are down regulated during gynoecium development when compared with SAM or leaf tissue. Cluster C3 (1060 genes) includes genes with putative roles in both, SAM and gynoecium development and including *ALCATRAZ (ALC), ARF5, DORNROSCHEN-LIKE (DRNL), HEC3, NTT, SQN, ULTRAPETALA* (*ULT*), *BEL1*, and *JAG*.

Cluster C8 includes 628 genes highly expressed in the leaf and gynoecium, but downregulated in the SAM, with CNA being the only known carpel regulator member. The cluster containing genes with SAM-only expression (C7) is surprisingly small with only 758 genes, as is the cluster C10 combining all 393 genes strongly expressed only in the last two stages of gynoecium development. Genes with high expression in the first two gynoecium development stages, less strong expression in leaf and SAM tissues are collected in cluster C9 (550 genes) and include *PIN1, PID, FIL, ANT, ETT, and ARF4*.

Next, we were interested in the biological processes reflected by the clusters and identified overrepresented GO terms (Fig. 3, Supplemental table 3). Cluster C1 including genes upregulated throughout gynoecium development shows enriched terms related to metabolic and transcriptional regulation, while the contrasting cluster C2 shows enriched terms related to general biosynthesis and metabolism. Metabolic processes are depleted in C3, a cluster similar to C1, but with the weakest expression in leaves. GO terms related to metabolic processes are also depleted in C6 that contains genes with the highest expression in S3 and S4, suggesting weaker metabolic activity during gynoecium development if compared to the SAM and leave tissues. In C6, terms related to fertilization and zygotic development are enriched. Cluster C9 includes genes highly expressed during S1 and S2 showing enriched terms related to the cell cycle and nucleic acid metabolism. Cluster C10 includes genes that are nearly exclusively expressed during S3 and S4, and shows enriched terms related to the import of nutrients, mainly sugars.

Next, we were interested to see which processes change during gynoecium development and how the gynoecium differs from leaf and SAM tissues. We compared co-expression clusters with contrasting patterns (Fig. 4), to elucidate the differences in enriched GO-terms between the set of genes expressed in the carpel when compared to other tissues as well as between the early (S1, S2) vs. late (S3, S4) carpel development (for the complete analysis see Supplemental Table X). GO terms related to hormone response are overrepresented only in late carpel development stages (Fig. 4). Cell cycle related genes are underrepresented genes in cluster C2 but highly overrepresented in genes upregulated in early stages of carpel development. Photosynthesis-related genes are overrepresented in C2 and in late stages of carpel development, while they are under-represented in cluster C1 and early carpel development. RNA-splicing related genes are overrepresented in cluster C1 and early carpel development suggesting that differential splicing may play a role in carpel morphogenesis. Genes involved in the regulation of gene expression are overrepresented throughout carpel development as are floral organ development genes.

**Figure 4:**
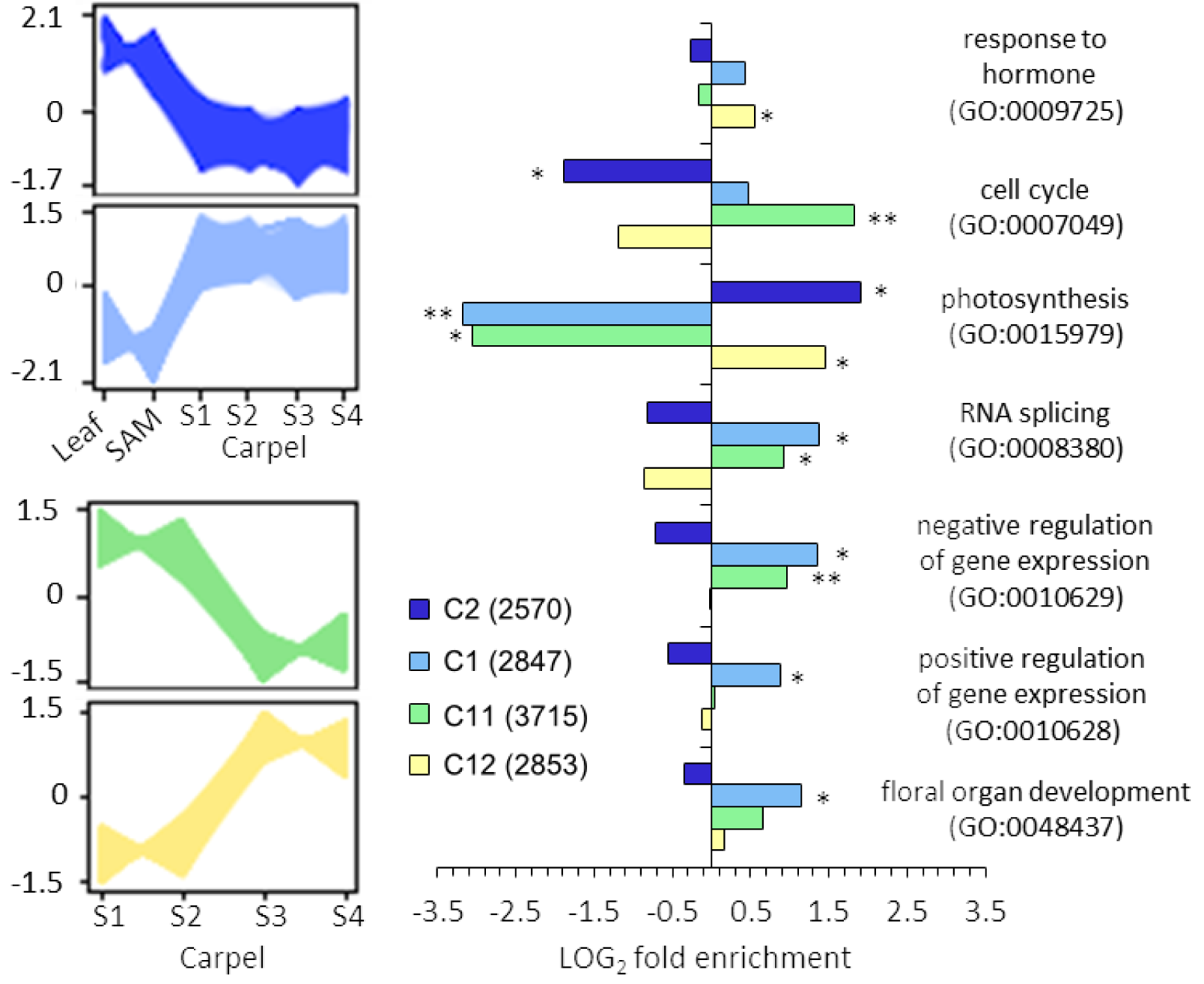
Over-representation of GO terms in clustered co-expression data. Co-expression clusters of the four carpel developmental stages with (blue clusters) and without (yellow and green clusters) expression data from SAM and leaf tissue with contrasting pattern are shown to the left. log2 fold enrichments of relevant GO-terms, representing larger sets of semantically similar terms, is shown on the right (* p-value < 0.005 – 10^−10^, ** p-value < 10^−10^).

In summary, our data show a succession of events, starting from upregulation of photosynthesis and downregulation of cell cycle activity in leaf and SAM. In early stages of carpel development, photosynthesis plays no major role but genes involved in cell cycle, regulation of gene expression and floral development are upregulated. In late stages, phytohormone response and photosynthesis-related genes are upregulated.

### Digital gene expression approaches can identify novel developmental regulators

Transcriptome analysis is a useful tool to clarify co-expression of gene clusters and single genes, but we were interested to know if it could also identify genes of hitherto unknown function that can be assigned as gynoecium developmental regulators. As proof of concept, seven genes with specific expression patterns were selected for reverse genetic analysis. While three SALK insertion lines showed no obvious fertility defects, four were significantly decreased in fertility (Fig. 5). These were named *LEF1-4* for *LESSER FERTILITY1-4* (Fig. 5A). *lef1* is mutated around 150 bp 5’ from the coding sequence of B3 domain family gene (AT5G46915) and has low expression restricted to S1 and S2 of gynoecium development. *lef2* has a insertion in the only exon of a DOF binding transcription factor encoding gene (AT5G66940) and, as *LEF1*, is restricted in its expression to the first two developmental stages. *LEF3* encodes an AP2/B3 transcription factor (AT3G17010) with high S1 and moderate S2 expression and the insertion is located in the second exon. *LEF4* also codes for an AP2/B3 transcription factor (AT3G46770) and strongly expressed in S2 and S3 and the insertion is in the first exon. The siliques of *lef1-4* are ranging from 9.4 −16.6% shorter and with 5.7 – 20.3 % fewer seeds than the wild type (Fig. 5C and D), all shown to be significantly different from Col-0. We were then interested if the LEF1-4 genes are integrated in the regulatory network shown in Fig. 2 and searched the upstream regions for transcription factor binding sites identified in ChIP-Seq experiments (Fig. 5E). Each gene is regulated by one MADS-box protein complex including AG and at least two MADS-box protein complexes bind to each promoter, suggesting that the LEF1-4 genes are under direct control of floral homeotic protein complexes.

**Figure 5:**
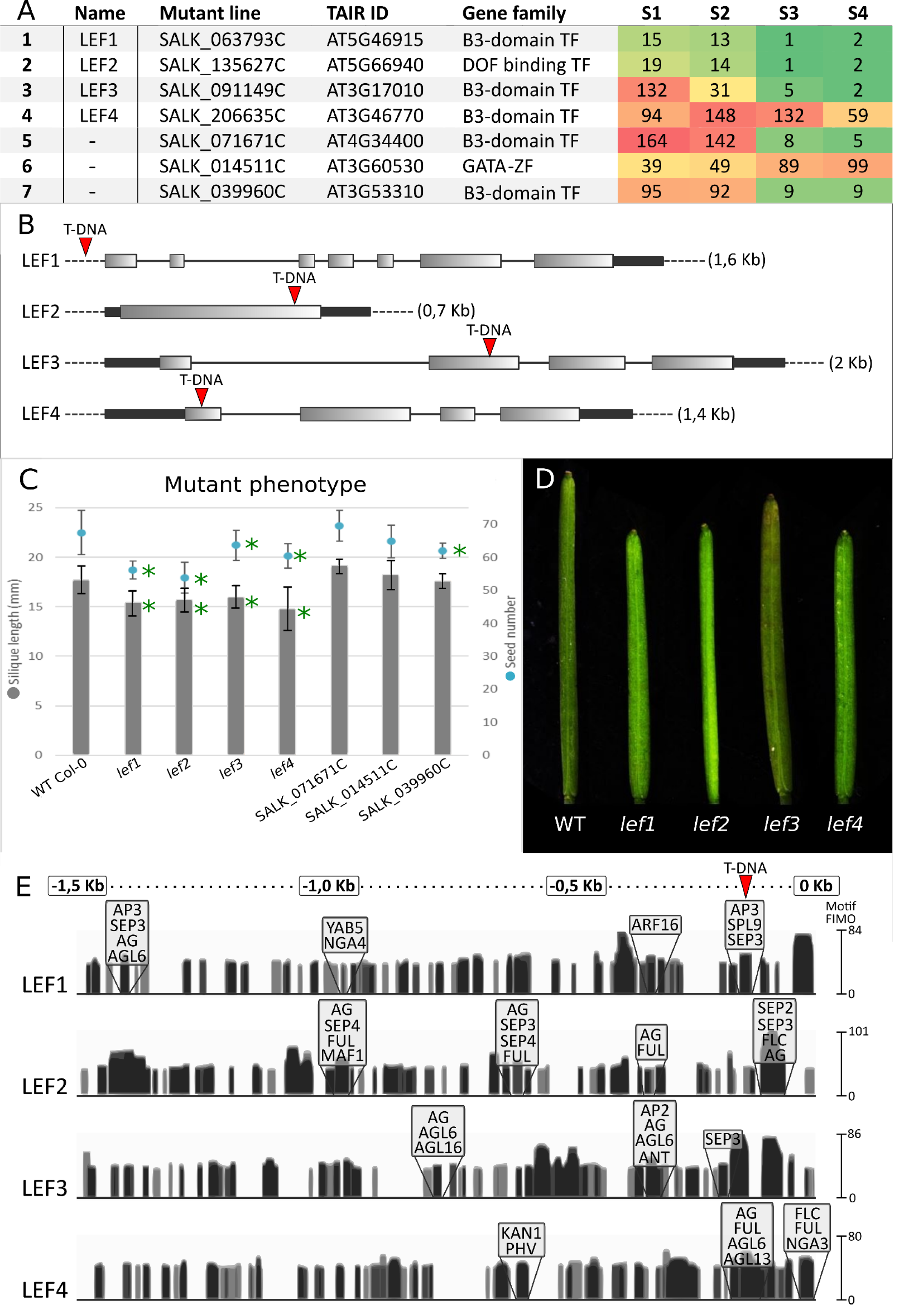
Candidate mutant analysis. A) An overview of the chosen genes and SALK insertion lines, gene families and expression. B) intron-exon structure of LEF-genes and locations of T-DNA insertion for each LEF-gene SALK-line C) Silique length and seed counts for all seven SALK lines (n=30 siliques). Given are the means and standard deviation. A star denotes significant difference of α > 0.05 to Col-0. D) Phenotypes of Col-0 wild type and the lef1-4 mutants showing significant defects in silique length and seed number E) LEF1-4 locus promoter analysis of TF binding verified by ChIP-Seq (http://www.chip-hub.org/). Y-axis labelling: FIMO (Find Individual Motif Occurences) is calculated by a log-likelihood ratio score for each position of the binding motif in the sequence database and includes q-values for each position by false discovery rate analysis. Binding sites for TRs 1,5 Kb upstream and highlighted genes with known regulatory functions in flower development. Areas with TF binding activity upstream the transcription start site are highlighted, bar height indicating the binding motif FIMO score (Grant et al., 2011) and colour tone indicating overlapping binding sites for multiple different binding entries. The red arrowhead indicates the site of the T-DNA insertion of the lef1 mutant.

## Discussion

Here, we use LMD RNAseq to generate expression data with high temporal resolution to resolve global transcriptional dynamics specific to gynoecium development. However, high specificity in transcriptome analysis may often go at the expense of sensitivity (probability to represent a particular transcript in the library), accuracy (how well the read quantification corresponds to actual mRNA concentration), and precision (technical variation of the quantification) (Ziegenhain et al., 2017). Here, we can show that transcription factor genes with known low levels of expression, such as *NGA2, NGA3, NGA4*, or *HEC1* (with RPKM of 14, 8, 7, and 1, respectively, Klepikova et al., 2016) are picked up by our approach and show TPM values of 29, 9, 2 and 6 (Supplemental table 1). These numbers compare well in magnitude with the RPKM values taken from Klepikova et al., (2016) demonstrating a high level of accuracy. Distance correlation analysis between biological replicates analysed in this study show that the transcriptomes of the two early and the two late stages are clearly distinct (Kivivirta et al., 2019). However, of the four LMD RNAseq replicates that were analysed for each stage, three clustered closely together and those were used for the analysis. Most likely, we have reached the morphological and genetic limit of differentiation between stages, and more fine-grained analysis by LMD would be sub optimal in terms of accuracy. Single-cell transcriptome analyses would be more suitable to e.g. identify transcripts of a specific, small-scaled tissue type, such as the *HEC1, 2, 3* genes which are expressed mainly in the few cells that later one will form the transmitting tract (Gremski et al., 2007), but their overall contribution to the transcriptome is very low (Fig. 1B). Single cell RNA sequencing (scRNA-seq) of developing gynoecia can improve sensitivity but relies in many cases on Fluorescence Assisted Cell Sorting (FACS). However, where fluorescent marker lines labelling specific tissues or cell types are limited, detection of cell types is difficult, and even more so for those cell types that form only a very small proportion in a tissue (Rich-Griffith et al., 2020). Here, this would apply to e.g. the dwindling stem cell population at early stages of gynoecium development or the placenta formation.

Also, the role of protein turnover on morphogenesis requires attention, when assessing the dynamics of transcriptional activity in developmentally active tissues. Stability varies among plant proteins, ranging from several hours to months with an average total protein half-life of 4-6 days (Li et al., 2017; Scheuerwater et al., 2000), but more specific data for TR’s turnover during developmental processes is not available. Thus, some transcriptional regulators may be active for a prolonged time even though their transcripts can no longer be detected. However, the effect of these stable proteins may be limited as it is diluted while the tissue increases in cell number. For example, *ANT, KNAT2, CRC, CUC3, NUB*, and *ETT* are required for CMM tissue differentiation but are mainly expressed at early stages and possibly, the proteins they encode persist for long. Moreover, while *HEC1, 2*, and *3* are expressed at very low levels throughout gynoecium development, their proteins may be particularly stable as their phenotypes are striking (Gremski et al., 2007; Schuster et al., 2015).

### High resolution transcriptome analyses reveal subfunctionalization between closely related homologs

The *SEP* genes are known for their importance in flower development and organ and meristem identity (Pelaz et al., 2000; Ditta et al., 2004) but so far, only little research has been published regarding each gene’s specific role in gynoecium development. *SEP* genes act partially redundantly in flower development, such that only the quadruple *sep1 sep2 sep3 sep4* mutant fails to form floral organs (Ditta et al., 2004), and *SEP3* is thought to be most important for floral organ identity as it forms most protein interactions with other MADS-box proteins (Immink et al., 2009). Moreover, it mediates ternary complex formation between AG and STK, AG and SHP1, AG and SHP2, SHP1 and SHP2, STK and SHP1, and STK and SHP2, all involved in carpel and ovule development (Favaro et al., 2003). However, our transcriptome analysis shows substantial differential dynamics of between the *SEP* genes (Fig. 1A), suggesting subfunctionalization of this gene family in gynoecium development. *SEP4* is generally expressed at a low level, but SEP1 and SEP2 are expressed strongly in the two early stages while *SEP3* is most strongly expressed in the two later stages. While the ternary complex formation of *SEP3* is well researched, the role of SEP1 and SEP2 has not been elucidated in much detail and they have fewer interactors among MADS proteins. Moreover, their ability for cooperative DNA binding differs between individual SEP proteins (Jetha et al., 2014). While the *sep1 sep2 sep3* mutant fails to form carpels (Pelaz et al., 2000), adding a single functional *SEP1* allele to the triple mutant restores carpel formation (Favaro et al., 2003). However, based on their strong expression during early carpel development we suggest important, but hitherto unknown roles for *SEP1* and *SEP2* in gynoecium development and a high degree of redundancy based on their sequence similarity and expression pattern, and possibly dimerization of SEP1 and SEP2 with non-MADS proteins. Severe subfunctionalization and extreme reduction in expression of *SEP* genes was also observed in several plant species, for example in *Gerbera hybrida*, whose genome includes seven *SEP* genes. While one of them, *GRCD6*, is hardly expressed, the other six genes diverge strongly in their expression pattern and function, and several distinct phenotypes were observed in the gynoecium when individual *SEP* genes were downregulated (Zhang et al., 2017). Gene duplication followed by subfunctionalization thus seems to be common to *SEP* homologs. The MADS-box genes *SHP1* and *SHP2* serve as second example for expression divergence of highly redundant genes. Neither of the single mutants displays a phenotype, but the double mutants are defective in dehiscence zone formation (Liljegren et al., 2000). Possibly, *SHP2* has an earlier function in gynoecium development as it expressed also in the early stages of gynoecium development and even stronger in late stages. In contrast, *SHP1* is hardly expressed in early stages and only moderately in the late stages (Fig. 1A). In addition, their interaction partners for dimerization differ, while SHP1 interacts with SEP3, SEP1, STK, AGL13, and AG, SHP1 interacts with SEP3, SEP1, and AGL6 only (Fig. 2A, Supplemental Table 3). However, SEP3 mediates interaction of SHP2 and STK as well as SHP2 and AG (Favaro et al., 2003), suggesting that subfunctionalization based on different dimerization partners overridden by ternary complex formation.

Similarly, *HEC* genes, known for their function in phytohormone control during gynoecium development (Schuster et al., 2015), show a peculiar pattern of expression. *HEC1* is expressed in the later stages especially at S3 where it interacts with SPT to control carpel fusion. The lesser known *HEC2* starts with strong early expression but completely ceases after S2 and *HEC3* is most expressed at the S2. The specific function of each HEC-gene is still mostly unclear but the transcriptomic data suggests a specific role for each of the three genes in carpel morphogenesis. Our data shows a replacement of early interaction of SPT-HEC2 with SPT and HEC1, HEC3, IND and ALC (Fig. 2A). Similar replacements can be observed in other hub proteins such as TPL and AG, which exchange interactions over time (Fig. 1A, B, Fig. 2A).

### Prediction of genetic interactions in gynoecium development

Negative or positive correlation of gene expression during gynoecium development can support predicted genetic interactions. For example, *SEU* and *LUG* together repress *AG* in the outer whorls of the flower (Franks et al., 2002) but we found strong expression of both of these genes also in the gynoecium with the highest expression during S1-3. This is in line with the *seu lug* phenotype in the gynoecium, characterized by lack of organ growth and carpel fusion (Franks et al., 2002) suggesting a continuous repression of hitherto unknown target genes during gynoecium development. Interestingly, both genes are also expressed significantly higher than *AG* in the gynoecium, suggesting that additional regulatory factors may be needed for the floral identity network regulation, protection of *AG* expression and proper gynoecium formation.

The protein interaction map (Fig. 2) provides a simplified overview of the regulatory dynamics during carpel development. Interactivity of the regulatory proteins is highly complex; however, relevant interactions are determined by presence and strength of expression at given time. Proteins like AG, PI, AP1, SEP1, SEP3, SHP1, SHP2 and STK interact with five or more regulatory partners each and expression of most of these proteins is established at carpel initiation or even before that. Our data suggests that interactions are at their highest complexity when tissue determinacy is established at the initiation of organogenesis as it has been described previously (Ò’Maoiléidigh et al., 2014). This can be observed as a high complexity of interactions during the initiation of carpel development (Fig. 2B).

### Co-expression clusters reveal temporal emphasis on gene expression regulation during gynoecium development

Comparing transcriptomes of four gynoecium stages, leaf and SAM tissues by co-expression clustering (Fig. 3) shows that that the majority of genes highly expressed throughout carpel development is at most weakly expressed in leaf and SAM tissue, suggesting a high level of difference between these tissues. Further, more co-upregulated genes are shared between the SAM and gynoecium than in leaf and gynoecium development suggesting closer similarity of gynoecium and SAM tissue. However, this may be due to rapid expansion of the organ combined with later arising meristematic activity from the carpel margins. The evolutionary ancestor of the carpel is thought to be leaf-like (Becker, 2020; Moubayidin and Ostergaard, 2017) and consequently, the transcriptional program of gynoecia should be more similar to that of leaves than the SAM. However, this might be an over simplified view as gene regulation related to photosynthesis is a major contribution to the leaf transcriptome but plays only a minor role in gynoecium development, as photosynthesis-related genes are enriched only in a single cluster comprising leaf and late gynoecium stages. With regard to developmental regulation, the leaf primordium is meristematic at its inception and, during growth, meristematic potential is restricted to the margins, reminiscent of different SAM zones (Alvarez et al., 2016). However, the leaf transcriptome does not reflect these spatially differentiated leaf tissue types or developmental stages. Interestingly, the observation that at/after stage 9 of *A. thaliana* flower development, gynoecium development shifts from bilateral to radial growth (Moubayidin and Ostergaard, 2017) is consistent with our data on TR expression (Fig. 1). Several transcriptional regulators change their expression between S2 and S3; for example, the adaxial/abaxial regulators *FIL, CRC*, and *KAN2* show a decline in expression after S2 suggesting that abaxial/adaxial polarity required for bilateral growth is established in S1 and S2 and subsequent radial growth requires different regulators. However, many regulatory processes seem to require maintenance throughout gynoecium development. For example, the C1 cluster includes genes upregulated throughout gynoecium development but not in leaf and SAM tissues and is most strongly enriched in regulators of gene expression, splicing and floral development.

Moreover, the shift of activity between S2 and S3 becomes also obvious when overrepresented GO terms are compared between genes upregulated in early (Fig. 4, C11) and late clusters (Fig. 4, C12). At the early stages, genes related to cell division, RNA splicing and regulation of gene expression enriched emphasizing the importance of early regulation of morphogenesis. In contrast, cluster C12 is enriched in photosynthesis related genes. This includes upregulation of genes required photosynthesis and for carbohydrate transport suggesting that the gynoecium may be a net sink organ but also contributes energy to the reproductive effort. Also, previous work has shown that flowers and fruits are not merely a cost to the carbon budget of the rest of the plant, but also contribute to this (Bazzaz et al., 1979; Gnan et al., 2017). Interestingly, in the case of gynoecium photosynthesis, developmental clues trigger upregulation of photosynthesis related genes and not light availability, because S4 is around 36 hours before anthesis (Smyth et al., 199o). Mizzotti et al., (2018) have shown that expression of photosynthesis, tetrapyrrol biosynthesis and plastid ribosomal proteins is strongest between three and six days after pollination and our data show that expression of many of these genes is already activated while the flower is still closed (Supplemental Table 3).

In summary, we describe a fine-scale map of transcriptional changes during gynoecium development as a resource to plant scientists. This provides a unique temporal perspective on global gene expression and protein complex formation potential suggesting a large number of new candidate developmental regulators orchestrating gynoecium development, four of which (LEF1-4) we have confirmed play a functional role.

## Materials and Methods

### Transcriptome assembly, heatmaps and interactome analysis

Raw sequencing reads of the four stages of *A. thaliana* gynoecium development (Kivivirta et al., 2019) were used to generate the transcriptomes (GenBank: Bioproject accession PRJNA549137). Trimming, quality testing, assembly and annotation were carried out in CLC workbench version 11.0.1. (QIAGEN, Hilden, Germany) as previously described in Kivivirta et al., 2019 with the the *A. thaliana* genome (Swarbreck et al., 2008). Gene expression heatmaps were constructed with Heatmapper: expression (Babicki et al., 2016). Euclidean distance of absolute values of gene expression were used with the expression values of a list of genes derived from Reyes-Olalde et al., 2013; Pfannebecker et al.; 2017a; Pfannebecker et al., 2017b; Parenicova et al. 2003 and Ojolo et al., 2018. Gene functions and families were based on earlier publications for each gene. The genes with expression of TPM <1 were left out of the analysis. For the complete list of genes, their functions and expression, see Supplemental Table 1. Protein-protein interaction maps were constructed with the GeneMANIA (Franz et al., 2018) app in cytoscape 3.8.0 (Shannon et al., 2003). Protein-protein interactions were searched for a set of carpel regulatory genes (Supplemental Table 1). Genes with no known interactions with other carpel regulatory genes were discarded. Information on gene families, change of gene expression and intensity of expression was added to the interaction map after the analysis. Change of binary logarithmic of expression was applied for Fig. 3A. Expression strengths for Fig. 3B and C is based on the absolute TPM values of gene expression.

### Co-expressed clusters

Automatically partitioned co-expression clusters were generated with Clust front-end version 1.0.0 (Abu-Jamous & Kelly 2018). The datasets were automatically normalised. Cluster tightness was set to 2 and minimum cluster size to 40 genes. Genes with flat expression were filtered out of the analysis. The SRA files additionally included in the analysis were SRR3581346 (SAM) and SRR3581838 (leaf blade) (Klepikova et al., 2016). GO enrichment for gene sets were analysed with PANTHER 15.0 gene ontology enrichment (Mi et al., 2019, GO version Mar 2020). The results of a Fisher’s Exact test using the *A. thaliana* reference list for overrepresentation analyses were corrected by calculating false discovery rates. The generated lists were then visualized using REViGO (Supek et al., 2011,) in the version available in June 2020 (GO version Jan 2017). For the visualization of redundant GO terms with REViGO, uncorrected p-values of overrepresentation tests were grouped based on their semantic similarity utilizing the inbuilt SimRel function with the *A. thaliana* reference and an allowed similarity setting of 0.7. The generated lists and treemaps were further processed with the DrasticData Treemapping tool (drasticdata.nl, Delft, The Netherlands) to achieve a better visualization.

### SALK mutant analysis

*A. thaliana* cv. Col-0 and the SALK mutant line plants were grown in peat, perlite mixture (3:1) in long day conditions. The mutants were self-pollinated to achieve homozygous insertion lines (Supplemental Table 4). SALK mutant lines were verified to be homozygous by genotyping the lines with locus- and insert-specific primers. Genotyping was done in two separate reactions with locus specific LbB + RP and LP + RP primers to verify presence of the specific insert and absence of the wild type locus (Supplemental Table 4). Siliques from 30 developing flowers were analysed for morphological abnormalities, gynoecium length and seed number at the stage of silique ripening (Smyth et al., 1990: stage 17) for each mutant line. Statistical significance was evaluated using the single factor ANOVA and t-test. The siliques were recorded with Leica DM550 (Leica Application Suite 4.3.0, Wetzlar, Germany) and the lengths of the siliques were measured with ImageJ (https://imagej.nih.gov/ij/). The intron-exon structures and protein binding site motifs for TRs upstream *LEF1-4* genes were based on ChIP-Hub (http://www.chip-hub.org). FIMO binding maps use Plant transcription factor database (http://planttfdb.cbi.pku.edu.cn/) as a source for protein binding sites.

## Supplemental material

**Supplemental table 1:** Detailed information on expression and function of genes related to gynoecium development (Fig. 1)

**Supplemental figure 1:** Heatmap of the MIKC type MADS-box genes. The heatmap figure illustrates the expression of MIKC type MADS-box genes at the four stages of development.

**Supplemental table 2:** Protein interactions of known developmental regulators. The table illustrates the experimentally verified physical protein interactions and the sources of information.

**Supplemental table 3:** Automatically partitioned Co-expressed clusters. The table illustrates the co-expression clusters, their contents, and the GO enrichment analysis.

**Supplemental table 4:** SALK-mutant analysis and genotyping. The table includes a summary of the candidate genes, SALK-mutant lines, genotyping primers, genotyping results and example pictures.

## Acknowledgements

We thank all members of the Becker lab for thoughtful discussion on the presented material. We are indebted to David Smyth for suggestions on the manuscript. Additionally, we thank the students Henri Hoffmann and Julian Garrecht for their help. We thank the German Research Foundation (DFG, project BE2547/14-1) for the main funding of this work. AB and SdF thank the DAAD-Conacyt collaborative grant numbers 267803 to SdF and 57273492 to AB.

## Author contribution

KK and AB designed the study and wrote the manuscript. KK performed digital gene expression analysis, mapping expression data to protein interaction data, and loss-of-function mutant analysis. CR conducted co-expression clusters and GO enrichment analyses. DH assembled the transcriptomes and calculated TPM values, and SdF and NMM analysed phytohormone-related genes. Figure preparation by KK and CR. All authors contributed, read and approved the final manuscript.

